# Visualizing Rev1 Catalyze Protein-template DNA Synthesis

**DOI:** 10.1101/2020.04.10.036236

**Authors:** Tyler M. Weaver, Luis M. Cortez, Thu H. Khoang, M. Todd Washington, Pratul Agarwal, Bret D. Freudenthal

## Abstract

During DNA replication, replicative DNA polymerases may encounter DNA lesions, which can stall replication forks. One way to prevent replication fork stalling is through the recruitment of specialized translesion synthesis (TLS) polymerases that have evolved to incorporate nucleotides opposite DNA lesions. Rev1 is a specialized TLS polymerase that bypasses abasic sites as well as minor-groove and exocyclic guanine adducts. It does this by using a unique protein-template mechanism in which the template base is flipped out of the DNA helix and the incoming dCTP hydrogen bonds with an arginine side chain. To observe Rev1 catalysis at the atomic level, we employed time-lapse X-ray crystallography. We found that Rev1 flips out the template base prior to binding the incoming nucleotide. Binding the incoming nucleotide changes the conformation of the DNA substrate to orient it for nucleotidyl transfer, and this is not coupled to large structural changes in the protein like those observed with other DNA polymerases. Moreover, we found that following nucleotide incorporation, Rev1 converts the pyrophosphate product to two mono-phosphates, which drives the reaction in the forward direction. Following nucleotide incorporation, the hydrogen bonds between the incorporated nucleotide and the arginine side chain are broken, but the templating base remains extrahelical. These post-catalytic changes prevent potentially mutagenic processive synthesis by Rev1 and facilitate dissociation of the DNA product from the enzyme.

## Introduction

Cells are tasked with efficiently and faithfully replicating the genome each round of cell division (1-3). During replication, the replicative DNA polymerases encounter a variety of DNA lesions that can stall the replication fork, thus promoting genomic instability (4, 5). One way the cell prevents this is through the recruitment of specialized translesion synthesis (TLS) polymerases that have evolved unique mechanistic and structural properties rendering them capable of bypassing DNA lesions that stall the replication fork (6, 7). Although both replicative and TLS polymerases perform the same general nucleotidyl transferase reaction, in which they extend a newly synthesized daughter strand of DNA, the mechanistic details of the reactions differ among the two polymerase types in order to provide the proper efficiency and fidelity necessary for each. In the case of a replicative DNA polymerase, the enzyme binds an incoming deoxynucleotide triphosphate (dNTP) and two associated metal ions, while sampling for proper Watson-Crick base pairing to the templating base. In most cases, dNTP binding is coupled to a polymerase conformational change in which the enzyme closes around the nascent base pair. Moreover, upon proper organization of the active site, the primer terminal 3′-OH undergoes deprotonation and in-line nucleophilic attack at Pα of the incoming dNTP resulting in the generation of a phosphodiester bond and a pyrophosphate (PP_i_) moiety (1-3, 8-11).

Rev1 is one of these specialized TLS polymerases with a preference for bypassing abasic sites as well as minor-groove and exocyclic guanine adducts (12-16). Rev1 predominantly inserts cytosine over all other dNTPs, which is advantageous during bypass of bulky adducted guanine residues (17-19). The first crystal structures of the Rev1 pre-catalytic ternary complex (Rev1/DNA/dCTP) revealed it uses a novel protein-templating mechanism (20). In this mechanism, Rev1 displaces the templating guanine into an extrahelical position. The templating guanine is subsequently replaced by an arginine side chain, which acts as a protein-template to hydrogen bond with the Watson-Crick face of the incoming dCTP (21). Additional structures of Rev1 in complex with a templating abasic site, γ-HOPdG, and BP-N2-dG have shown Rev1 uses a similar mechanism to bypass multiple types of DNA damage (12, 22, 23).

Several unanswered questions regarding the Rev1 reaction mechanism hamper our understanding of how it catalyzes this unusual means of DNA synthesis. To this point, the protein template Rev1 uses for DNA synthesis requires the templating base be evicted from the active site, however, whether eviction of the templating base occurs concurrently with incoming dCTP binding is unclear. Additionally, the post-catalytic steps remain unknown at the atomic level. This includes how the dCMP disengages from the templating arginine and whether the templating base remains extrahelical after dCMP insertion. Finally, a molecular-level mechanism that limits Rev1 from performing multiple mutagenic cytosine insertions during TLS has not been deciphered. To address these outstanding questions, we have used a combination of time-lapse X-ray crystallography, molecular dynamic simulations, and biochemical approaches.

## Results

To observe the Rev1 catalytic cycle at the atomic level, we employed time-lapse X-ray crystallography (Fig. 1A) (24-29). To accomplish this, we first generate binary Rev1/DNA complex crystals with a templating guanine. Importantly, Rev1 interacts with a templating guanine and damaged guanines in an identical manner (12, 22, 23). These binary Rev1/DNA crystals are subsequently transferred into a cryoprotectant solution containing dCTP and CaCl_2_ to generate Rev1 ground-state ternary (Rev1/DNA/dCTP) complexes. Importantly, Ca^2+^ facilitates nucleotide binding but does not support catalysis (24-29). To initiate catalysis, the Rev1 ground-state ternary complex crystals are subsequently transferred to a cryoprotectant solution containing either MgCl_2_ or MnCl_2_. The crystals are then flash-frozen at various time points prior to collecting X-ray diffraction data (Fig. 1A). The resulting structures provide high-resolution snapshots of the Rev1/DNA complex during active site organization, catalysis, and post-catalytic events.

**Figure 1.**
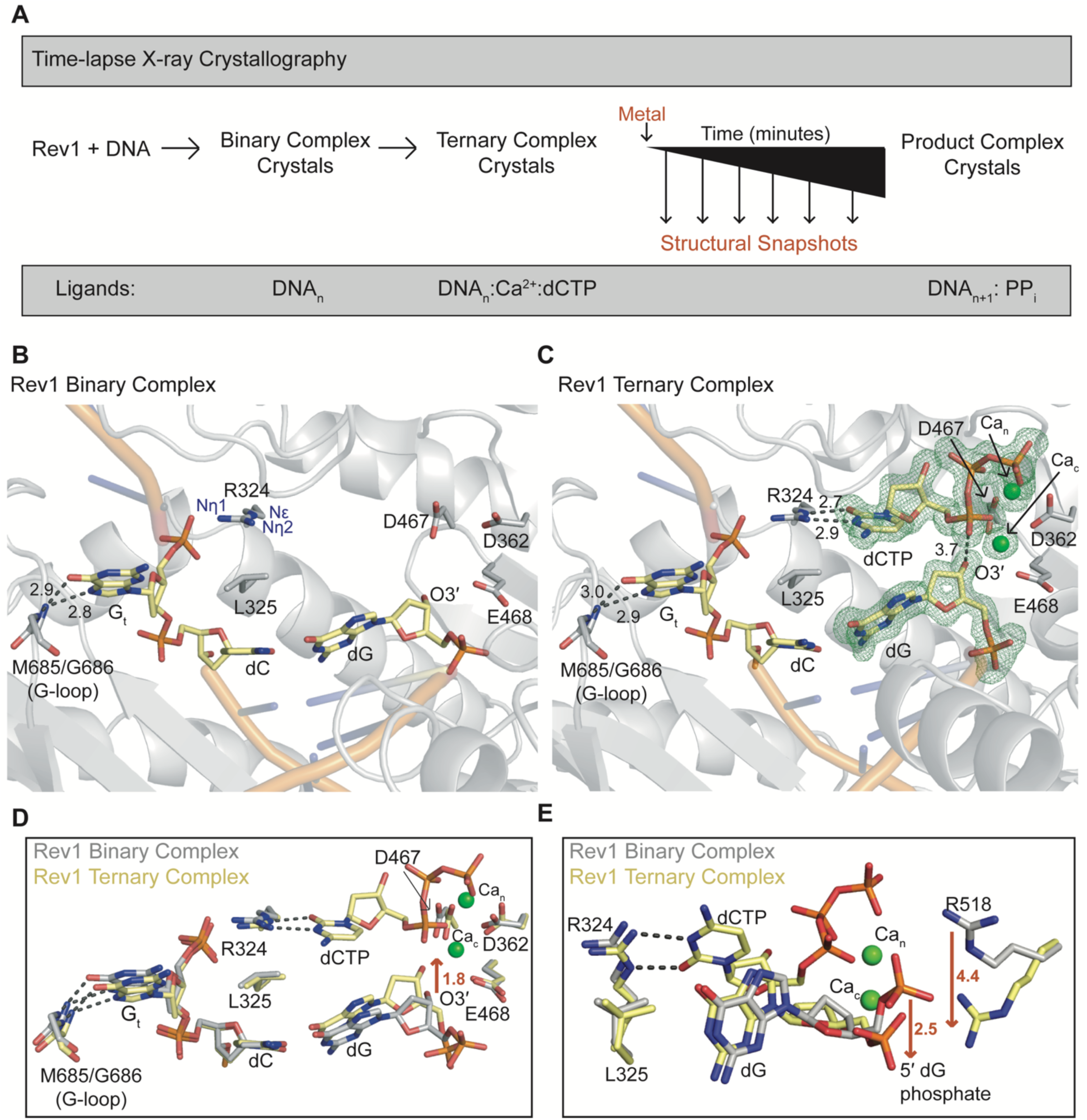
Rev1 binary and ternary crystal structures. **(A)** The time-lapse X-ray crystallography approach. The ligands bound to each Rev1 state are indicated at the bottom. An active site closeup of the Rev1 **(B)** binary and **(C)** ternary crystal structures. The nucleic acid residues are shown in yellow and Rev1 in grey. A Fo-Fc OMIT map OMIT map contoured at σ=3.0 around the two active-site calcium ions, incoming dCTP, and primer terminal dG is shown as a green mesh. **(D and E)** An overlay of the Rev1 binary (grey sticks) and ternary (yellow sticks) complexes are shown in two orientations to indicate the movement at the primer termini, phosphate backbone, and R518. Key protein and DNA residues are indicated in each panel.

### Pre-catalytic organization of the Rev1 active site

To decipher the steps necessary for organization of the Rev1 active site, we solved a 2.05 Å structure of the Rev1 binary complex in space group P2_1_2_1_2_1_ (Table 1). To our knowledge, this is the first Rev1 binary structure and it shows the templating guanine (G_t_) is evicted from the active site and displaced ∼90° from the DNA helix (Fig. 1B). The evicted G_t_ is stabilized within a hydrophobic pocket and forms hydrogen bonds between N7 and O6 of its Hoogsteen edge and the backbone amides of M685 and G686 in the Rev1 G-loop (Fig. 1B). The void generated by the evicted G_t_ is occupied by residues R324 and L325 of the Rev1 N-digit helix (Fig. 1B). L325 has been hypothesized to be important for evicting and keeping the templating G_t_ out of the DNA duplex, while R324 serves as the protein-template for binding the incoming dCTP through the Nε and Nη of the guanidinium group (Fig. 1B). Displacement of the G_t_ and the positioning of R324 as the protein-template is the first step in organizing the Rev1 active site for binding of the incoming dNTP.

**Table 1.** Data collection and refinement statistics for Rev1 crystal structures.

|  | Binary | Ternary | Intermediate | Product<br>Two<br>phosphates | Product<br>One<br>phosphate |
| --- | --- | --- | --- | --- | --- |
| **Data collection** |  |  |  |  |  |
| Space group | P2<sub>1</sub>2<sub>1</sub>2<sub>1</sub> | P2<sub>1</sub>2<sub>1</sub>2<sub>1</sub> | P2<sub>1</sub>2<sub>1</sub>2<sub>1</sub> | P2<sub>1</sub>2<sub>1</sub>2<sub>1</sub> | P2<sub>1</sub>2<sub>1</sub>2<sub>1</sub> |
| Cell dimensions |  |  |  |  |  |
| *a*, *b*, *c* (Å) | 62.2, 73.9, 117.6 | 62.3, 73.9, 117.3 | 62.3, 73.7, 117.2 | 62.4, 73.8, 117.1 | 62.2, 73.5, 117.3 |
| $\alpha$, $\beta$, $\gamma$ (°) | 90, 90, 90 | 90, 90, 90 | 90, 90, 90 | 90, 90, 90 | 90, 90, 90 |
| Resolution (Å) | 25 – 2.05 | 50 – 1.40 | 25 – 1.8 | 50 – 2.20 | 25 – 2.05 |
| *R*<sub>meas</sub><sup>a</sup> (%) | 0.147 (0.473) | 0.070 (>1) | 0.104 (>1) | 0.115 (0.690) | 0.089 (0.772) |
| *I*/σ*I* | 10.3 (2.7) | 25.4 (1.06) | 15.3 (1.05) | 8.0 (1.32) | 11.7 (1.4) |
| *cc*1/2<sup>b</sup> | 0.644 | 0.426 | 0.494 | 0.435 | 0.444 |
| Completeness<sup>a</sup> (%) | 99.6 (97.0) | 99.5 (95.1) | 99.4 (97.7) | 99.6 (98.9) | 99.7 (98.8) |
| Redundancy<sup>a</sup> | 3.1 (2.9) | 4.0 (2.5) | 4.2 (3.1) | 3.9 (2.3) | 4.3 (2.5) |
| **Refinement** |  |  |  |  |  |
| Resolution (Å) | 22.4 – 2.05 | 35.2 – 1.40 | 24.68 – 1.80 | 24.77 – 2.20 | 24.96 – 2.05 |
| No. reflections | 104907 | 198840 | 95877 | 111434 | 148755 |
| *R*<sub>work</sub>/*R*<sub>free</sub> | 20.5/25.8 | 18.94/21.22 | 18.17/23.43 | 18.53/25.82 | 17.76/25.12 |
| No. atoms |  |  |  |  |  |
| Protein | 3518 | 3540 | 3565 | 3534 | 3539 |
| DNA | 571 | 571 | 571 | 571 | 591 |
| Water | 340 | 730 | 568 | 150 | 179 |
| B-factors (Å<sup>2</sup>) |  |  |  |  |  |
| Protein | 24.58 | 19.99 | 25.60 | 40.09 | 32.81 |
| DNA | 34.62 | 23.66 | 32.54 | 48.14 | 41.40 |
| Water | 31.80 | 32.43 | 36.55 | 40.74 | 35.82 |
| R.m.s deviations |  |  |  |  |  |
| Bond length (Å) | 0.008 | 0.007 | 0.007 | 0.009 | 0.008 |
| Bond angles (°) | 0.931 | 0.935 | 0.840 | 1.030 | 1.006 |
| PDB ID |  |  |  |  |  |

<sup>a</sup> Highest resolution shell is shown in parenthesis

**Table 1 continued.** Data collection and refinement statistics for Rev1 crystal structures
|  | Product<br>Zero<br>phosphates | WT Product<br>MnCl<sub>2</sub> soak | L325G<br>Product complex | R518A<br>Ternary complex |
| --- | --- | --- | --- | --- |
| **Data collection** |  |  |  |  |
| Space group | P2<sub>1</sub>2<sub>1</sub>2<sub>1</sub> | P2<sub>1</sub>2<sub>1</sub>2<sub>1</sub> | P2<sub>1</sub>2<sub>1</sub>2<sub>1</sub> | P2<sub>1</sub>2<sub>1</sub>2<sub>1</sub> |
| Cell dimensions<br>a, b, c (Å) | 62.3,72.4,118.49 | 62.0,73.8,117.6 | 61.96,73.6,117.6 | 62.25,73.77,116.81 |
| α, β, γ (°) | 90,90,90 | 90,90,90 | 90,90,90 | 90,90,90 |
| Resolution (Å) | 25 – 1.70 | 50 – 1.98 | 50 – 2.55 | 25 – 1.64 |
| *R*<sub>meas</sub><sup>a</sup> (%) | 0.091 (>1) | 0.163 (>1) | 0.165 (0.818) | 0.128 (0.834) |
| *I*/*σ**I* | 27.5 (0.8) | 11.8 (1.2) | 7.8 (1.2) | 8.4 (0.58) |
| *cc*1/2<sup>b</sup> | 0.548 | 0.630 | 0.649 | 0.616 |
| Completeness<sup>a</sup> (%) | 99.9 (99.5) | 100 (100) | 99.4 (98.3) | 99.8 (99.2) |
| Redundancy<sup>a</sup> | 9.0 (2.8) | 7.9 (5.3) | 3.8 (2.9) | 4.1 (3.0) |
| **Refinement** |  |  |  |  |
| Resolution (Å) | 24.5 – 1.70 | 42.68 – 1.98 | 35.1 – 2.55 | 24.89 – 1.64 |
| No. reflections | 113949 | 61018 | 31944 | 49632 |
| *R*<sub>work</sub>/*R*<sub>free</sub> | 19.62/23.11 | 20.51/26.74 | 20.17/28.45 | 19.70/24.42 |
| No. atoms |  |  |  |  |
| Protein | 3535 | 3483 | 3479 | 6989 |
| DNA | 591 | 591 | 591 | 886 |
| Water | 418 | 258 | 60 | 455 |
| B-factors (Å<sup>2</sup>) |  |  |  |  |
| Protein | 28.61 | 29.06 | 43.51 | 20.79 |
| DNA | 34.37 | 40.64 | 52.19 | 33.83 |
| Water | 37.67 | 34.28 | 41.55 | 28.28 |
| R.m.s deviations |  |  |  |  |
| Bond length (Å) | 0.006 | 0.015 | 0.009 | 0.019 |
| Bond angles (°) | 0.845 | 1.605 | 1.115 | 1.502 |
| PDB ID |  |  |  |  |

To capture the ground-state ternary complex of Rev1 with an incoming dCTP, binary complex crystals were soaked in a cryoprotectant containing 5 mM dCTP and 50 mM CaCl_2_ (Fig. 1A). The resulting crystal diffracted to 1.40 Å and is in the P2_1_2_1_2_1_ space group (Table 1). In the ternary complex, the G_t_ remains in an identical conformation as observed for the binary complex (Fig. 1C). The incoming dCTP forms a planar hydrogen bonding interaction with the Nε and Nη of the R324 side chain via the O2 and N3 of the dCTP Watson-Crick edge (Fig. 1C). Structural superimposition of the binary and ternary complexes reveal the Rev1 active site does not undergo large conformational changes upon dCTP binding (Fig. 1D). Only subtle ∼1.0 Å changes at residues D362 and D467 are observed upon nucleotide binding and metal coordination. In contrast, substantial structural changes occur upon dCTP binding at the primer terminus. The 5′ phosphate backbone of the primer terminal dG is shifted 2.5 Å compared to the position in the binary state (Fig. 1E). The R518 side chain, which coordinates the non-bridging oxygen of the 5′ dG phosphate backbone, shifts 4.4 Å to track the primer strand movement (Fig. 1E). This shift in the primer strand results in a 1.8 Å shift of the primer terminal O3′ towards the Pα of the incoming dCTP (Fig. 1D). These movements result in the O3′ of the primer terminus being 3.7 Å from Pα of the incoming dCTP and in proper orientation for in-line nucleophilic attack. Together, the Rev1 binary and ternary complex structures indicate that movement at the primer terminus is important for pre-catalytic organization of the Rev1 active site.

### Characterizing Rev1 catalyze protein-template DNA synthesis

To capture structural snapshots during catalysis, Rev1/DNA/dCTP ternary complex crystals were transferred to a second cryoprotectant lacking dCTP and containing MgCl_2_ instead of CaCl_2_ for 5 min (Fig. 1A). The resulting crystal diffracted to 1.80 Å and was in space group P2_1_2_1_2_1_ (Table 1). This intermediate structural snapshot contained both reactant (dCTP) and product (dCMP and PP_i_) states, as evident by electron density between Pα and Pβ as well as O3′ and Pα, respectively (Fig. 2A). Occupancy refinement determined the reaction is ∼30% product and ∼70% reactant. The position of active site residues and metal ions remain identical to the ground state ternary complex, thus indicating the Rev1 protein does not undergo large structural changes during catalysis. In contrast, significant movement occurs in the primer DNA strand that coincides with phosphodiester bond formation. The 5’ phosphate backbone of dG, formerly the primer terminal nucleotide, is in two conformations. The product conformation is shifted 4.0 Å proximal from the conformation in the reactant conformation (Fig 2B). This coincides with a 5.4 Å shift of R518, which coordinates the 5’ backbone phosphate of the dG (Fig 2B). The inserted dCMP in the product state shifts toward the primer terminal base pair by 1.1 Å compared to the incoming dCTP in the ground state. This shift results in loss of planarity between the dCMP and R324 (Fig 2B). The change in conformation of the inserted dCMP after bond formation likely disrupts the hydrogen bonding interaction between R324 and the Watson-Crick face of the inserted dCMP.

**Figure 2.**
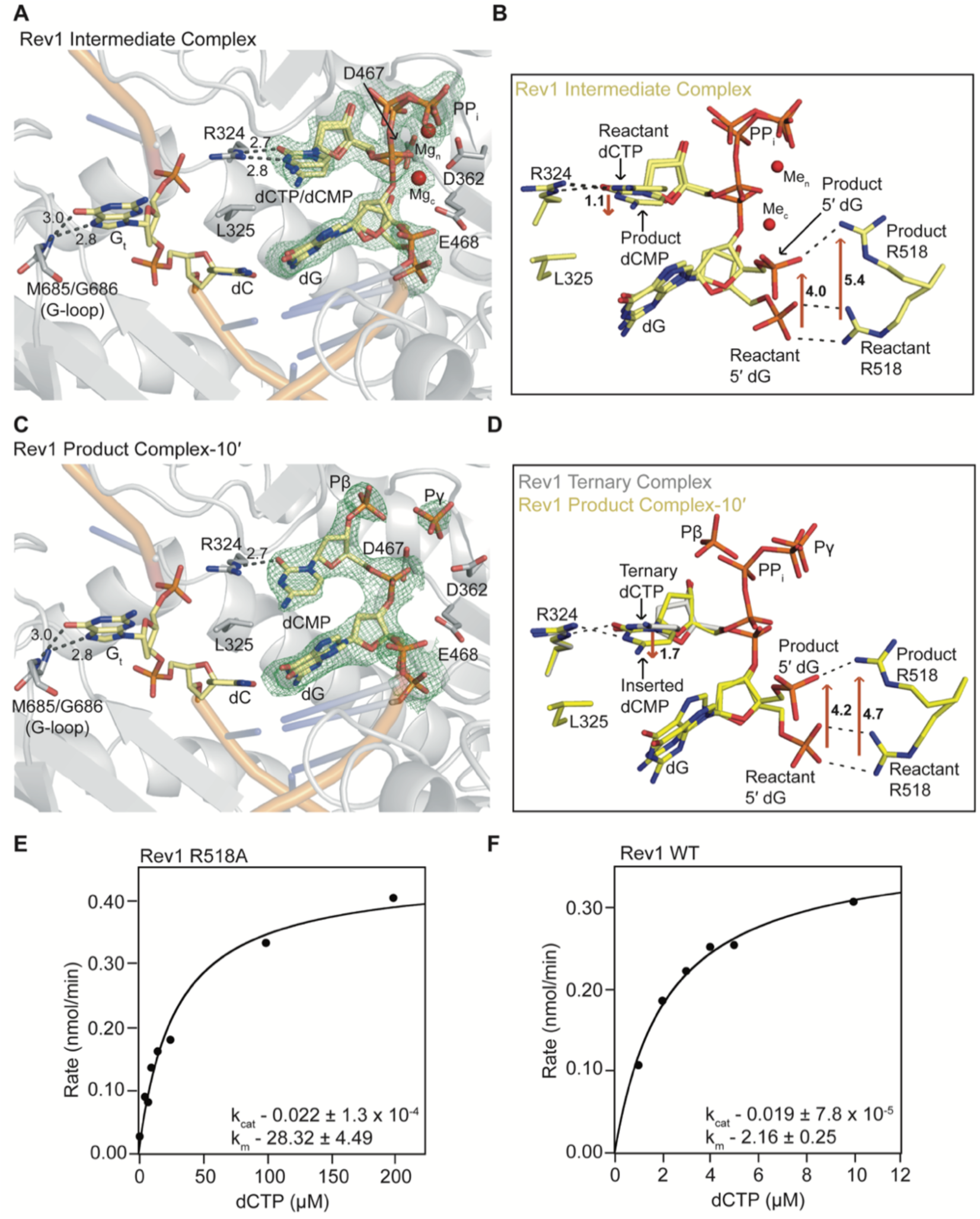
Rev1 protein-templated DNA synthesis. **(A)** An active site closeup of the Rev1 intermediate crystal structure. The nucleic acid residues are shown in yellow and Rev1 in grey. A Fo-Fc OMIT map contoured at σ=3.0 is shown as a green mesh. **(B)** A focused view of the structural movements in the Rev1 intermediate complex. The reactant and product states are indicated. The red arrows and distances (Å) highlight significant conformational differences. **(C)** An active site closeup of the Rev1 product complex after a 10’ soak in MgCl_2_. A Fo-Fc OMIT map OMIT map contoured at σ=3.0 is shown as a green mesh. **(D)** A focused view of the active site of the product complex (yellow sticks) with an overlay of the dCTP from the Rev1 ternary complex (grey sticks) shown for reference. Red arrows and distances (Å) are indicated to highlight significant movements. Plots of the steady state kinetics for **(E)** R518A and **(F)** WT Rev1 are shown with the k_cat_ and K_m_ indicated for each graph. Steady state kinetics were performed in triplicate.

To obtain a product complex, ternary complex crystals (Rev1/DNA/dCTP) were soaked in a cryoprotectant containing MgCl_2_ for 10 min (Fig. 1A). The resulting crystal diffracted to 2.20 Å and shows the nucleotidyl transfer reaction is complete (Table 1 and Fig. 2C). Consistent with the prior structural snapshots, the structural changes observed post catalysis are isolated to the primer DNA strand and not the Rev1 protein. The inserted dCMP shifts 1.7 Å towards what was the primer terminal dG, which results in a loss of planarity between the dCMP and R324 (Fig. 2D). The distances between R324 and the Watson-crick edge of the inserted dCMP are now 2.9 Å and 3.0 Å. This combined with the loss in planarity indicates the hydrogen bonding interaction between dCMP and R324 is lost or weakened (Fig. 2D). In addition, the dCMP is more dynamic in the product complex with a B-factor of 43.07 Å^2^ compared to 12.40 Å^2^ for the incoming dCTP in the ground-state ternary complex. The primer DNA strand also undergoes additional movement during product formation. The 5′ backbone phosphate of the dG, formerly the primer terminal nucleotide, exists in two conformations. Conformation one is similar to the conformation in the ground state ternary complex. In the second conformation, the dGMP 5′ backbone phosphate has moved 4.2 Å proximal to the position in the ternary complex (Fig. 2D). Similar to the transition from binary to ternary complex, the two conformations of R518 track the two conformations of the the dGMP 5′ backbone phosphate.

To understand the role of R518 in Rev1 catalysis, kinetic parameters were obtained for the insertion of dCTP by both the R518A variant and wild-type (WT) Rev1 protein. The steady-state kinetic analysis revealed Rev1 R518A has a k_cat_ of 0.022 +/- 1.3×10^−4^ min, which is almost identical to Rev1 WT at 0.019 +/- 7.8×10^−5^ min (Fig. 2E and F). In contrast, Rev1 R518A has a ∼15-fold lower K_m_ (28.32 +/- 4.49 μM) than Rev1 WT (2.16 +/- 0.25 μM) indicating that R518 is important for incoming dCTP binding (Fig. 2E and F). To understand the potential role of R518A in nucleotide binding, a ground-state ternary structure of Rev1 R518A with an incoming dCTP was solved to 1.64 Å (Table 1). The overall structure of WT and R518A Rev1 in the ternary ground state structures is very similar with an RMSD (Cα) of 0.109. In addition, the Rev1 active site including the conformation of R324, L325, the incoming dCTP, and metal coordinating residues is identical between the two structures (Fig. S1). However, B-factor analysis revealed that the O3′ of the primer terminus in the R518A ternary ground-state structure was more dynamic at 17.26 when compared to the O3′ in the WT ternary ground state structure at 13.92. This is likely due to disruption of the hydrogen bonding between R518 and the non-bridging oxygen of the 5′ dG phosphate backbone in the alanine mutant Rev1 (Fig. S1B). Together, these data are consistent with R518 playing in important role in stabilizing movement of the primer terminus to organize the active site for incoming nucleotide binding.

### MD Simulations of Rev1

To gain insight into the dynamics within the Rev1 active site during the catalytic cycle we employed computational modeling of the Rev1 binary, ternary ground-state, and product structures. These molecular dynamic (MD) simulations were performed for 1 microsecond (1 µs) and provide insight into the interactions of the evicted G_t_, incoming dCTP, and inserted dCMP. The X-ray crystal structures show the evicted G_t_ is stabilized via hydrogen bonds between N7 and O6 of the Hoogsteen edge with the backbone amides of M685 and G686, respectively (Fig. 1 and 2). By monitoring these hydrogen bonding distances throughout the simulation, we can gain insight into the stability of the extrahelical G_t_. During the binary simulation, these interactions are broken around 220 ns of the MD simulation as indicated by the increased distance between M685_n_ -N7 and G686_n_ -O6 (black line, Fig. 3A). This results in the G_t_ returning back to the intrahelical orientation within the DNA helix. In contrast, the G_t_ remained extrahelical for the entire MD simulation of the ternary (dCTP bound) and product (dCMP inserted) complexes, as indicated by the constant hydrogen bonding distances between M685_n_ -N7 and G686_n_ -O6 (red and green lines, Fig. 3A). Together these simulations indicate the templating G_t_ remains extrahelical upon binding and insertion of the incoming dCTP.

**Figure 3.**
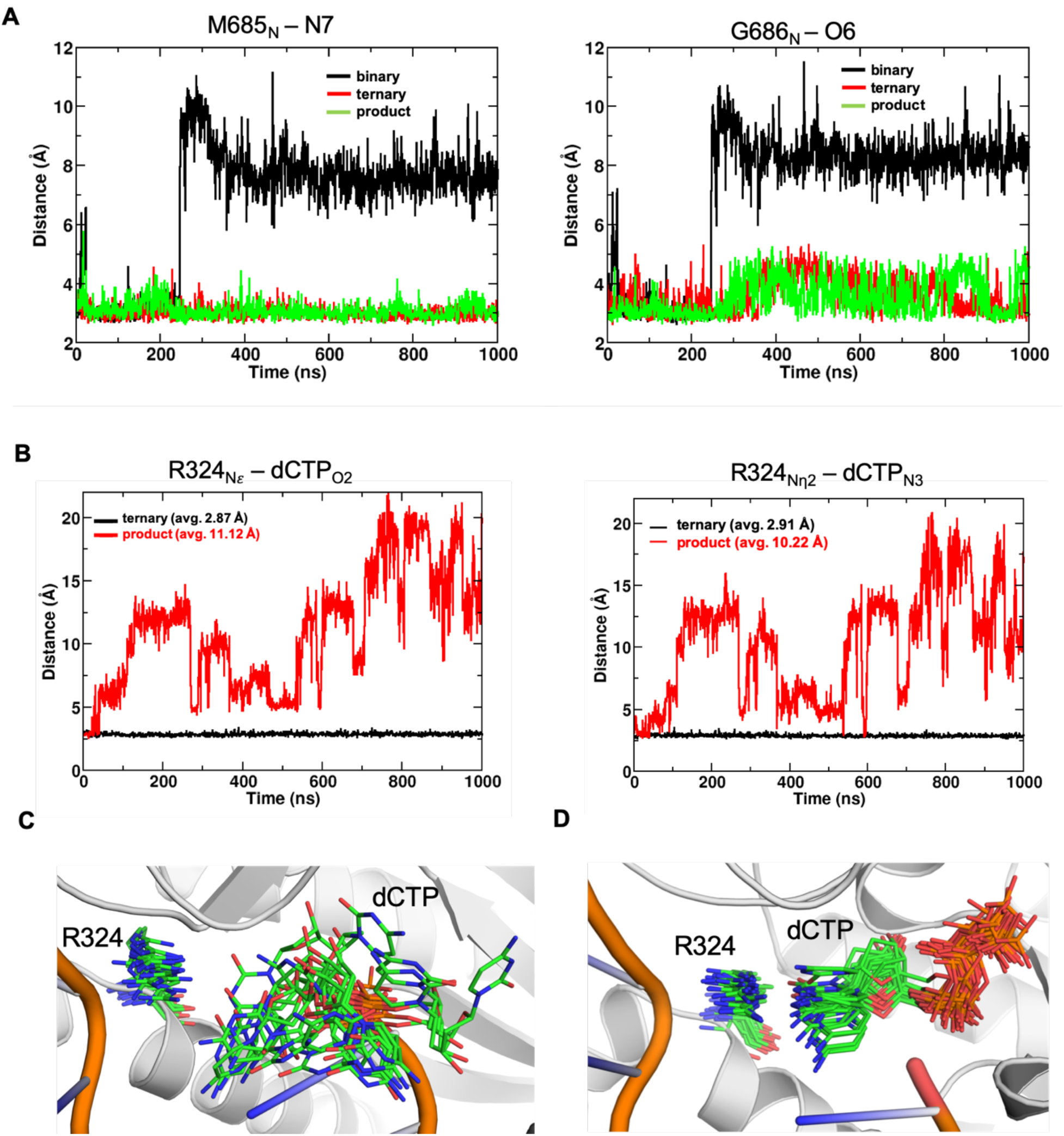
Molecular dynamic simulations of Rev1 binary, ternary, and product complexes. **(A)** Distance profiles for the M685_N_ – N7 (left panel) and the G686_N_ -O6 (right panel) in the Rev1 binary (black line), ternary (red line) and product (green line) MD simulations. This behavior was reproduced by a second independent trajectory in which the interactions break after about 200 ns. **(B)** Distance profiles for the R324_NE_ and dCTP_O2_ (left panel) and the R324_Nn2_ and dCTP_n3_ (right panel) in the Rev1 ternary (black line) and product (red line) MD simulations. The average distance between these atoms is indicated for the entire trajectory. **(C)** 20 conformations from MD simulations for the product complex (each 50 ns apart) showing the conformation of dCTP and R324. **(D)** 20 conformations from MD simulations for the ternary complex (each 50 ns apart) showing the conformation of dCTP and R324.

To characterize the dynamics of the incoming dCTP and inserted dCMP we monitored the distances between the Ne and Nn2 of R324 and O2 and N3 of the dCTP, respectively, for the Rev1 ternary ground-state and product complex. The ternary ground-state MD simulation indicates R324 interacts strongly with the incoming dCTP, while in the product complex MD simulation the inserted dCMP is dynamic and breaks the structural contacts with R324. In the ternary ground-state complex, short and stabilizing hydrogen bonding interactions (<3 Å) were observed to be present between R324 and the incoming dCTP for the entire 1 µs simulation (Fig. 3B, black line). In contrast, the interactions between the inserted dCMP and R324 were broken immediately at the start of the simulation of the product complex (Fig. 3B, red line). In the product complex, the inserted dCMP is very dynamic and adopts multiple orientations with conformations sampling both in and out of the DNA helix (Fig. 3C), while in the case of ternary ground-state complex the structural interactions appear to keep the dCTP bound in a single conformation and hydrogen bonding to R324 (Fig. 3D). This result is consistent with the inserted dCMP breaking the hydrogen bonding interaction with R324 post-catalysis. Of note, throughout the product complex simulation the inserted dCMP does not hydrogen bond to the evicted G_t_.

### PPi breakdown in the Rev1 active site

The canonical DNA polymerase reaction results in phosphodiester bond formation and the generation of pyrophosphate (PP_i_). Interestingly, we did not observe PP_i_ in the Rev1 product complex and instead observed density for two monophosphates corresponding to Pβ and Pγ (Fig. 2C). This indicates hydrolysis of pyrophosphate occurred in the Rev1 active site following dCMP insertion, which is consistent with recent time-resolved crystallography with DNA Pol IV (30). The two monophosphates in the Rev1 active site are no longer bound by a nucleotide associated metal and subtle structural changes occur to alleviate the electrostatic repulsion from the negatively charged species within the Rev1 active site. The Pβ has shifted 2.7 Å away from D362 and D467, while remaining coordinated to R408, S402, and N414. The Pγ conformation is similar to the ground-state ternary complex, however D362 has shifted 1.7 Å and rotated 45° away from Pγ in the product structure (Fig. 4A). To confirm Rev1 hydrolyzes pyrophosphate in solution, we performed a primer extension assay while monitoring the amount of pyrophosphate or monophosphate generated by Rev1. The primer extension reactions resulted in ∼30-fold more monophosphate (10.89 ± 1.12 μM) than pyrophosphate (0.35 ± 0.15 μM), consistent with Rev1 mediated pyrophosphate hydrolysis during dCTP insertion (Fig. 4B).

**Figure 4.**
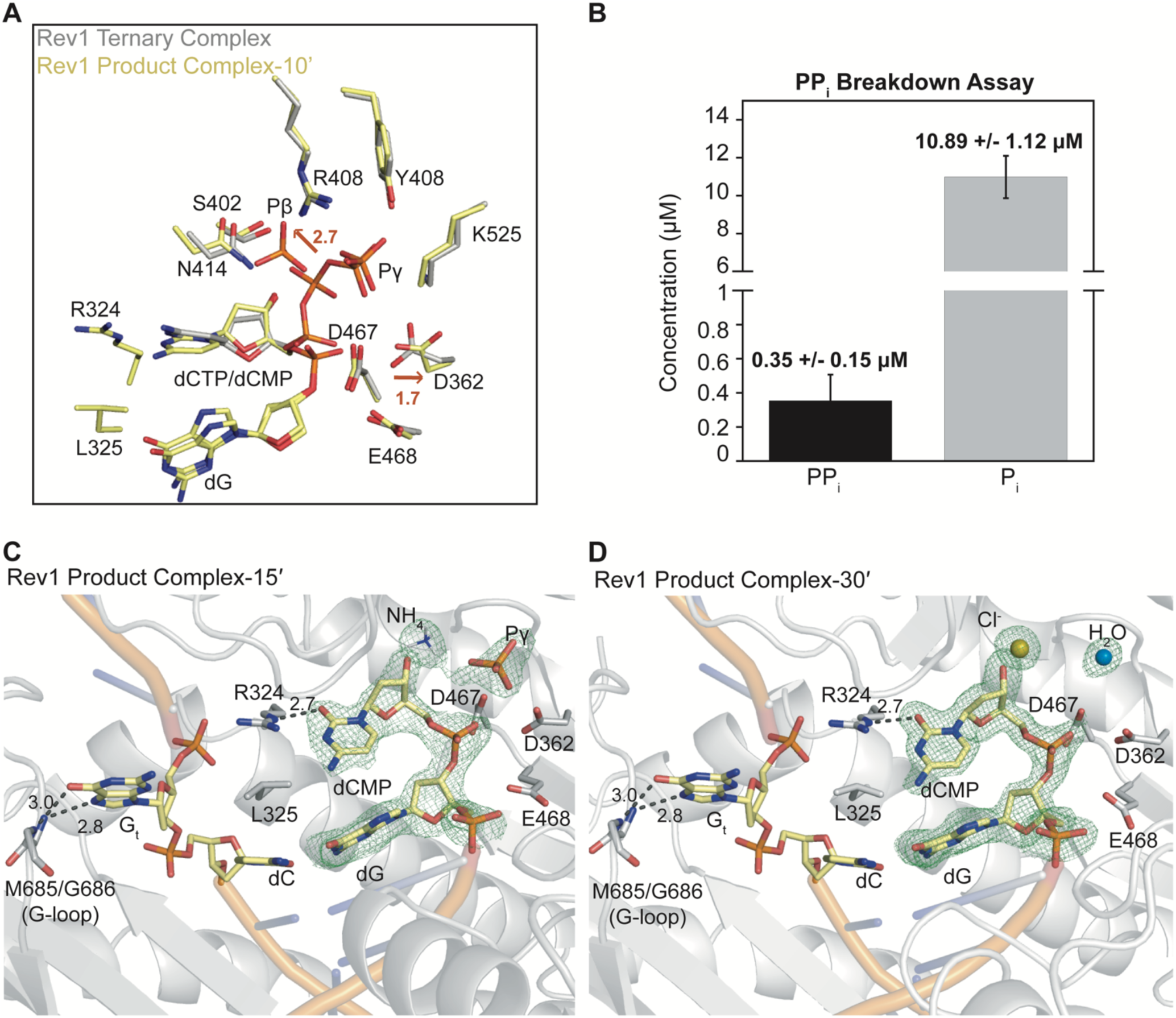
PPi breakdown and disassociation in the Rev1 active site. **(A)** A focused view of the PP_i_ movements in the active site of the product complex after a 10’ soak (yellow sticks) with an overlay of the Rev1 ternary complex (grey sticks) shown for reference. Red arrows and distances (Å) are indicated to highlight significant differences between the two structures. **(B) T**he concentrations of pyrophosphate (PP_i_) and monophosphate (P_i_) generated in solution during the Rev1 primer extension assay. Average of 3 replicate experiments. An active site closeup of the Rev1 product complex after a **(C)** 15’ and **(D)** 30’ soak in MgCl_2_. The nucleic acid residues are shown in yellow and Rev1 in grey. A Fo-Fc OMIT map contoured at σ=3.0 is shown as a green mesh. Key residues are indicated.

To determine the events that occur following pyrophosphate hydrolysis, two additional structures were determined after 15 and 30 min soaks in MgCl_2_. After 15 min, the Rev1/DNA product structure diffracted to 2.05 Å and was in space group P2_1_2_1_2_1_ (Table 1). In this structure, the Pβ has disassociated from the active site and the Pγ remains bound (Fig. 4C). An ammonium ion from the crystallization solution binds in the Pβ binding site after dissociation. After a 30 min soak in MgCl_2_, the Rev1/DNA product structure diffracted to 1.70 Å and was in space group P2_1_2_1_2_1_ (Table 1). In this structure, the Pγ has dissociated from the active site and is replaced by a water molecule, while a Cl^-^ ion is bound in the Pβ binding site (Fig. 4D). These structures indicate Rev1 utilizes a stepwise release of monophosphates with Pβ dissociating prior to Pγ. After monophosphate dissociation, the active site residues of Rev1 are in the same conformation as the binary (Rev1/DNA) structure, indicating that Rev1 has returned to the pre-nucleotide binding state. Despite phosphodiester bond formation, hydrolysis of pyrophosphate and release of monophosphates from the active site, the G_t_ remains extrahelical and has not base paired with the inserted dCMP (Fig. 4D).

### Rev1 processivity

The three Rev1/DNA product structures indicate the G_t_ remains extra-helical after catalysis has occurred. This suggests Rev1 may be unable to register shift or translocate to the next base to continue DNA synthesis with additional cytosine insertions. To better understand Rev1 processivity in solution, a primer extension assay was designed. In this assay, Rev1 is pre-incubated with a fluorescein-labeled DNA substrate and the reaction is then initiated with MgCl_2_ and 150-fold excess of non-labeled trap DNA. This reaction allows for Rev1 to continue primer extension until it dissociates from the DNA and is sequestered from its substrate by the trap DNA. In the presence of MgCl_2_ Rev1 largely extends the primer DNA strand by only a single nucleotide before it dissociates. Though a second, less efficient, insertion event is seen at higher dCTP concentrations (Fig. 5A). Manganese is often used as an alternative metal co-factor when studying DNA polymerase mechanism, as it enhances both nucleotide binding and catalysis. Therefore, to determine if additional insertion of the second nucleotide could be achieved, the primer extension assay was performed using MnCl_2_. In the presence of Mn^2+^, the processivity assay revealed insertion of one or two dCMPs before dissociation of Rev1 from the DNA substrate (Fig. 5B). Importantly, the second dCMP insertion by Rev1 is substantially more efficient in the presence of Mn^2+^ than Mg^2+^.

**Figure 5.**
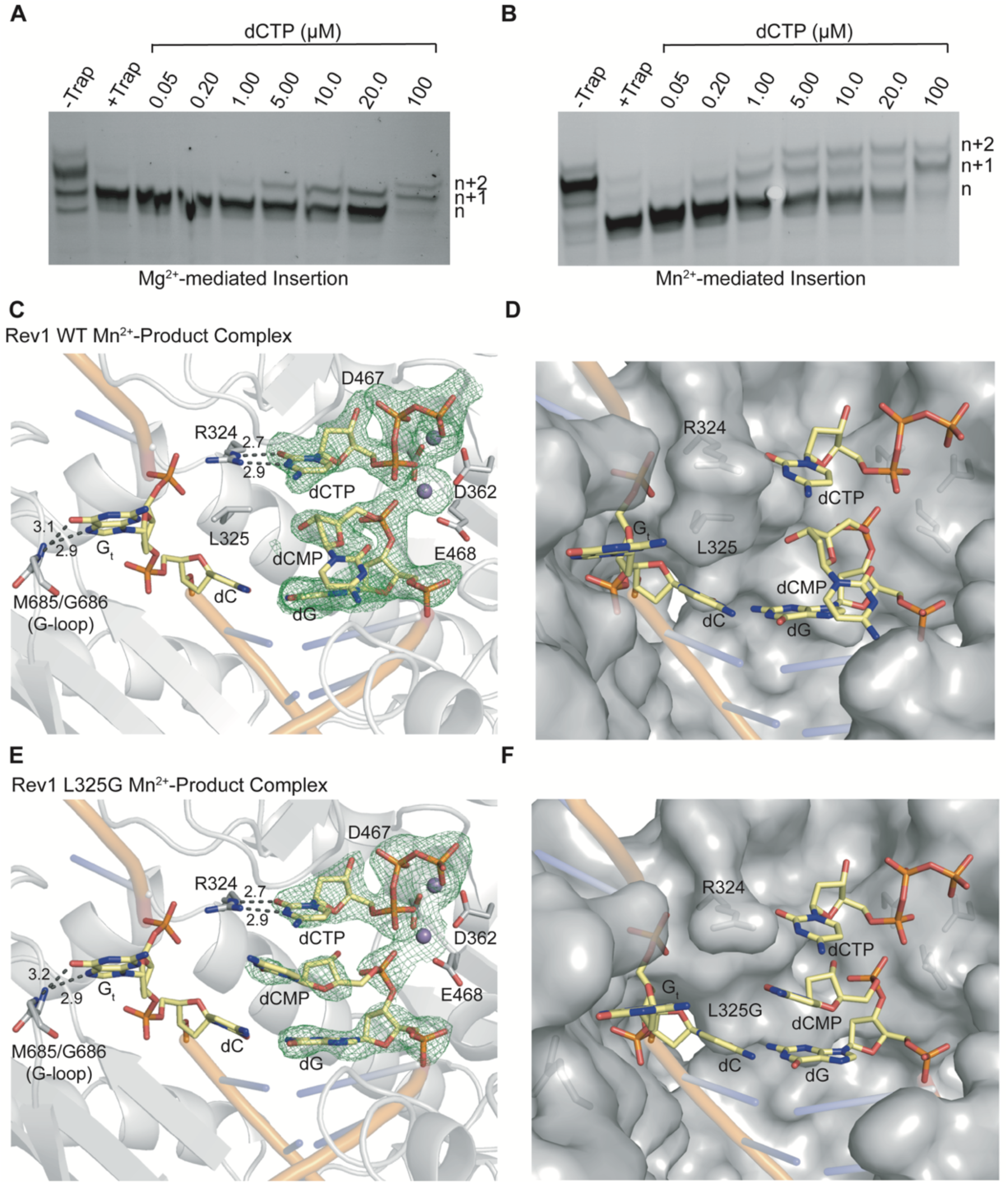
Rev1 processivity. The Rev1 processivity assay in the presence of MgCl_2_ **(A)** or MnCl_2_ **(B)**. Each processivity assay was repeated at least 3 times. **(C)** An active site closeup of the Rev1 product complex after a 240’ soak in MnCl_2_. The nucleic acid residues are shown in yellow, Mn^2+^ in purple, and Rev1 in grey. A Fo-Fc OMIT map contoured at σ=3.0 is shown as a green mesh. The inserted dCMP, second bound dCTP, and key residues are indicated. **(D)** The same structure and view as in panel C with Rev1 shown in a grey surface representation. **(E)** An active site closeup of the L325G Rev1 product complex after a 20’ soak in MnCl_2_. The nucleic acid residues are shown in yellow, Mn^2+^ in purple, and Rev1 in grey. A Fo-Fc OMIT map contoured at σ=3.0 is shown as a green mesh. The inserted dCMP, second bound dCTP, and key residues are indicated. **(F)** The same structure and view as in panel E with Rev1 shown in a grey surface representation.

To capture a structural snapshot of Rev1 inserting a second nucleotide, Rev1/DNA ternary complex crystals were soaked in a cryoprotectant containing MnCl_2_ for 240 min. The resulting crystal diffracted to 1.98 Å and was in space group P2_1_2_1_2_1_ (Table 1). In the Mn^2+^-mediated product structure, Rev1 has inserted a dCMP and bound a second incoming dCTP. The inserted dCMP, now at the n-1 position, has been displaced into the major grove to accommodate a second bound dCTP within the active site (Fig. 5C and 5D). Despite insertion of a dCMP, and the binding of a second dCTP, the evicted G_t_ remains extra helical. To determine if L325 prevents the Gt from base pairing with the inserted dCMP, we crystallized the L325G Rev1 binary complex. These crystals were transferred to a cryoprotectant containing dCTP and MnCl_2_ for 20 min. The resulting crystal diffracted to 2.55 Å and was in space group P2_1_2_1_2_1_ (Table 1). This structure showed a single dCMP has been inserted and a second dCTP bound in the active site of Rev1, similar to the WT Rev1 MnCl_2_ product complex (Fig. 5E). The void generated by the L325G mutation is occupied by the inserted dCMP and the G_t_ remains extra-helical (Fig. 5F). Together, this indicates that reincorporation of the G_t_ and base pairing with the dCMP likely requires dissociation of Rev1 from the DNA substrate and poses a barrier to Rev1 processivity.

## Discussion

Here, we utilized time lapse X-ray crystallography to obtain snapshots of Rev1 during catalysis. We identified movements of the DNA primer terminus that are important for both active site organization before catalysis and dissassocation of the products after catalysis. In addition, we find Rev1 hydrolyzes pyrophosphate after phosphodiester bond formation and releases the monophosphates from the active site in a stepwise manner. These structural snapshots also provide insight into the mechanism by which Rev1 prevents processive nucleotide incorporation to avoid aberrant insertions of cytosines after lesion bypass. Together, this work provides a fundamental advance in our understanding of how Rev1 catalyzes protein-template-directed DNA synthesis.

### Organization of substrates and products within the Rev1 active site

DNA polymerases bind the DNA and incoming nucleotide substrates and arrange them such that the 3*′*-oxygen of the primer termini and the α-phosphate of the incoming nucleotide are properly oriented for the nucleotidyl transfer reaction(1-3, 8-11). This occurs in two sequential steps: DNA binding and subsequent dNTP binding. In the case of most DNA polymerases, the DNA-binding step involves the DNA substrate binding to an open form of the polymerase. The duplex region of the DNA is approximately B-form, and the template base is positioned helically, *i*.*e*., it is positioned such that it would form a normal helical Watson-Crick base pair if the correct nucleotide were to bind opposite it. Our structure of the Rev1/DNA binary complex shows that Rev1 differs from other DNA polymerases in the DNA binding step. DNA binding by Rev1 involves displacement of the G_t_ from the DNA helix with R324 and L325 filling the space in the helix vacated by the flipped-out G_t_. Importantly, the position of R324 in the binary complex is poised for dCTP binding and is analogous to the position of the templating DNA base in the structures of other polymerase/DNA binary complexes.

In the case of most DNA polymerases, the nucleotide-binding step couples the association of the nucleotide and metal ions to conformational changes in the polymerases. Typically, this involves an open-to-closed transition of the fingers subdomain around the incoming nucleotide and the template base. These large conformational changes facilitate the organization of the DNA and incoming nucleotide substrates for optimal catalysis. Our structure of the Rev1 ternary complex shows that Rev1 does not use such a concerted mechanism to organize substrates within its active site. No large changes in the structure of the protein are observed upon incoming nucleotide binding. Instead there is substantial movement of the primer DNA strand to reposition the primer terminal O3*′* for an in-line nucleophilic attack on the α-phosphate of the incoming nucleotide. Despite the unique mechanism Rev1 uses for nucleotide binding, the resulting active site orientation for in-line nucleophilic attack on the α-phosphate is identical to other DNA polymerases.

In the product complex of other DNA polymerases, the incorporated nucleotide is base paired with the templating base(1-3, 8-11). The base-paired product is then released from the active site through re-opening of the fingers subdomain of the polymerase. Our structure of the Rev1 product complex shows that Rev1 does not do this. The G_t_ remains extrahelical in the Rev1 product complex, and the incorporated dCMP residue has shift by about 1 Å. This results in the loss of planarity between the incorporated dCMP and the side chain of R324 and the weakening of the hydrogen bonds between the C and R324. This suggests that Rev1 must dissociate from the DNA before the incorporated C can base pair with the template G. Furthermore, this dissociation may be facilitated by the movement of the primer terminus after catalysis that weakens the hydrogen bonds between R324 and the incorporated C.

### The role of pyrophosphate hydrolysis in polymerase function

The first evidence of pyrophosphate hydrolysis in the active site of a DNA polymerase was recently described for DNA Pol IV (30). Our data indicates Rev1 also breakdowns pyrophosphate in the active site *in crystallo* and in solution. Pyrophosphate hydrolysis in the active site of a DNA polymerase is proposed to make the DNA synthesis reaction energetically favorable (30). In the case of Rev1, it is likely important for driving the forward reaction in two ways. First, it makes phosphodiester bond formation energetically favorable without the need for a pyrophosphatase enzyme, as was hypothesized for DNA Pol IV. Second, it prevents the reverse reaction of pyrophosphorolysis, which can remove an added nucleotide.

Interestingly, the third metal seen in the product state of other polymerases studied by time-lapse crystallography has been proposed to play a role in facilitating catalysis, stabilizing the product state, and preventing the reverse reaction (24-29). Therefore, the breakdown of pyrophosphate by Rev1 could explain why a third metal is not seen during Rev1-mediated insertion. This is consistent with DNA Pol IV, which also did not contain a third metal in the product state for Mg^2+^-facilitated insertion(30). However, it is possible that pyrophosphate hydrolysis leads to a short-lived state containing the third product metal that we were unable to capture in the structural snapshots presented here. Ultimately, hydrolyzing pyrophosphate after phosphodiester bond formation would likely commit a DNA polymerase to the forward DNA synthesis reaction.

### Impaired processivity prevents aberrant cytosine insertions by Rev1

TLS polymerases have substantially lower fidelity than any other family of DNA polymerases. However, the low fidelity is balanced by the ability to bypass several types of DNA lesions that prove difficult for replicative polymerases to handle. Rev1 is an extreme example of a low-fidelity polymerase as it preferentially inserts cytosines regardless of the templating base. Until now, the mechanism that prevents extensive mutagenic dCMP insertions once Rev1 is recruited to DNA lesions was unclear. Our crystal structures of the Rev1/DNA product state reveal the G_t_ remains displaced even after catalysis occurs, indicating that translocation to the next base is unable to occur. Consistently, Rev1 largely inserts only a single nucleotide in the presence of Mg^2+^ in solution before dissociating from the DNA. Although a second dCMP insertion in the presence of Mn^2+^ is observed in solution, our crystallographic data shows Rev1 accomplishes this through reorganization of the primer DNA strand for binding a second dCTP without reincorporation of the G_t_ into the DNA helix. Together, this data suggests Rev1 must dissociate from the DNA substrate, instead of ratcheting or translocating like classical DNA polymerases, before additional aberrant cytosine insertions can occur. After dissociation, Rev1 may re-bind the DNA substrate, or more likely, be replaced by or recruit a DNA polymerase with higher fidelity and less mutagenic potential such as Pol ζ for continued DNA synthesis.

## Methods

### Preparation of DNA

For X-ray crystallography, 5′-ATC-GCT-ACC-ACA-CCC-CT-3′ (template strand) and 5′-GGG-GTG-TGG-TAG-3′ (primer strand) oligonucleotides were annealed 1x TE (10 mM Tris-8.0 and 1mM EDTA) by heating to 90 °C for 5 minutes before cooling to 4 °C using a linear gradient (−1 °C min^-1^). To generate the fluorescein-labeled DNA substrate for steady state kinetics and the Rev1 processivity assay, the 5′-GTA-CCC-GGG-GAT-CCG-TAC-GCC-GCA-TCA-GCT-GCA-G-3′ (template strand) and 5′-FAM-CTG-CAG-CTG-ATG-CGG-3′ (primer strand) oligonucleotides were annealed in 1x TE by heating to 90 °C for 5 minutes, cooling to 65 °C for 5 minutes before cooling to 10 °C using a linear gradient (−1 °C min^-1^). To generate the unlabeled trap DNA for processivity assays and the pyrophosphate breakdown assays, the 5′-GTA-CCC-GGG-GAT-CCG-TAC-GCC-GCA-TCA-GCT-GCA-G-3′ and 5′-CTG-CAG-CTG-ATG-CGG-3′ oligonucleotides in 1x TE by annealed by heating to 90 °C for 5 minutes, cooling to 65 °C for 5 minutes before cooling to 4 °C using linear gradient (−1 °C min^-1^). All DNA substrates were purchased from Integrated DNA Technologies.

### Cloning, Expression and Purification of Rev1

The yeast Rev1 construct was purchased from GenScript. All Rev1 proteins were expressed in BL21(DE3) plysS E. coli cells (Invitrogen). Cells were grown at 37 °C until an OD_600_ ∼0.7 was reached and protein expression induced with 0.1 mM IPTG at 20 °C overnight. Cell pellets were frozen and stored at -20 °C. For lysis, Rev1 cell pellets were resuspended in a lysis buffer containing 50 mM HEPES (pH-7.4), 150 mM NaCl, 1 mM EDTA, 1 mM DTT and a cocktail of protease inhibitors. The cells were lysed via sonication, lysate was cleared at 24,000 xg for one hours and the supernatant incubated with glutathione agarose resin (Goldbio) for 2 hours. The protein was washed on the glutathione beads with a high salt buffer containing 50 mM HEPES and 1M NaCl and protein cleaved overnight using PreScission Protease. The cleaved Rev1 protein was purified by cation-exchange chromatography using a POROS HS column (GE Health Sciences) and eluted from the column using a linear gradient of a buffer containing 1 M NaCl and 50 mM HEPES (pH-7.4). Rev1 was further purified by gel filtration using a HiPrep 16/60 Sephacryl S-200 HR (GE Health Sciences) in a buffer containing 250 mM NaCl and 50 mM Tris (pH-8.0). The purified Rev1 protein was frozen and stored at -80 °C. All protein concentrations were determined by absorbance at 280 nM using a NanoDrop One UV–Vis Spectrophotometer (Thermo Scientific).

### X-ray Crystallography

For X-ray crystallography, 5′-ATC-GCT-ACC-ACA-CCC-CT-3′ (template strand) and 5′-GGG-GTG-TGG-TAG-3′ (primer strand) oligonucleotides were used. The annealed DNA substrate (1.15 mM) was mixed with Rev1 protein (5-6 mg ml^-1^) and the Rev1-DNA binary crytals obtained in a condition with the reservoir containing 15-23% PEG3350 and 200 mM ammonium nitrate using the sitting-drop vapor diffusion method. The Rev1-DNA binary crystals were transferred to a cryoprotectant containing reservoir solution and 25% glycerol. Data were collected at 100 K on a Rigaku MicroMax-007 HF rotating anode diffractometer equipped with a Dectris Pilatus3R 200K-A detector system at a wavelength of 1.54 Å. Initial models were generated by molecular replacement using a previously solved Rev1/DNA ternary complex (PDB 5WM1) as a reference structure. Subsequent refinement was carried out using PHENIX and model building with Coot. All structure figures were generated using PyMOL (Schrödinger LLC).

### Steady State Kinetics

The primer extension reactions for steady state kinetic analysis were performed using a fluorescein-labeled oligonucleotide 5′-GTA-CCC-GGG-GAT-CCG-TAC-GCC-GCA-TCA-GCT-GCA-G-3′ (template strand) and 5′-FAM-CTG-CAG-CTG-ATG-CGG-3′ (primer strand). The reactions were performed using 10 nM Rev1 protein and 100 nM DNA substrate in a buffer containing 25 mM Tris (pH-8.0), 100 mM KCl, 5 mM MgCl_2_, 1 mM DTT and 100 μg mL^-1^ BSA. Initiation of the reactions were done by addition of dCTP at varying concentrations (WT-1, 2, 3, 4, 5 and 10 μM and R518A-1, 5, 7.5, 10, 15, 25, 100 and 200 μM) for 15 minutes and quenched using a loading dye containing 80% formamide 100 mM EDTA, 0.25 mg ml^-1^ bromophenol blue and 0.25 mg ml^-1^ xylene cyanol. Samples were heated to 95 °C and subsequently run on a 22% denaturing polyacrylamide gel. The gel was imaged using a Typhoon FLA 9500 imager (GE Health Sciences) and data processed in ImageJ. The steady state kinetics curves were fit using the Michalis-Menten equation:

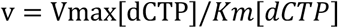

### Pyrophosphate Breakdown Assay

The primer extension reactions for pyrophosphate breakdown used oligonucleotides 5′-GTA-CCC-GGG-GAT-CCG-TAC-GCC-GCA-TCA-GCT-GCA-G-3′ and 5′-CTG-CAG-CTG-ATG-CGG-3′. Rev1 primer extension reactions were performed using 250 nM Rev1 and 15 μM DNA substrate in a buffer containing 25 mM Tris (pH-8.0), 5 mM DTT, 100 μM EDTA and 10% glycerol. The reactions were initiated by addition of 5 mM MgCl_2_ and 1 mM dCTP for one hour before quenching with 20mM EDTA. For pyrophosphate detection, the quenched reactions were incubated with a fluorogenic pyrophosphate sensor (MAK168, Sigma Aldrich) for 30 minutes. The fluorescence intensity was measured (λex =316 nm and λem = 456 nm) using a Horiba FluoroMax-4 Fluorimeter and the concentration of pyrophosphate in the samples were determined from a standard curve generated from using purified pyrophosphate. For monophosphate detection, the reactions were incubated with Malachite green dye (MAK308, Sigma Aldrich) for 30 minutes. The absorbance of the samples was collected at 620 nM using a NanoDrop One UV– Vis Spectrophotometer (Thermo Scientific) and the concentration of monophosphate determined using a standard curve generated from purified monophosphate.

### Rev1 Processivity Assay

The primer extension reactions for Rev1 processivity analysis were performed using a fluorescein-labeled oligonucleotide 5′-GTA-CCC-GGG-GAT-CCG-TAC-GCC-GCA-TCA-GCT-GCA-G-3′ (template strand) and 5′-FAM-CTG-CAG-CTG-ATG-CGG-3′ (primer strand). The reactions were carried out using 100 nM Rev1, 100 nM DNA substrate and increasing concentrations of dCTP(.05-100 μM) in a buffer containing 25 mM Tris (pH-8.0), 5mM DTT, 100 μM EDTA and 10% glycerol. The reactions were initiated using 5mM Mg^2+^ or 5 mM Mn^2+^, 15mM spermidine and 15 nM trap DNA made from oligonucleotides 5′-GTA-CCC-GGG-GAT-CCG-TAC-GCC-GCA-TCA-GCT-GCA-G-3′ and 5′-CTG-CAG-CTG-ATG-CGG-3. The reactions were quenched using a loading dye containing 80% formamide 100 mM EDTA, 0.25 mg ml^-1^ bromophenol blue and 0.25 mg ml^-1^ xylene cyanol. Samples were heated to 95 °C and subsequently run on a 22% denaturing polyacrylamide gel and the gel imaged using a Typhoon FLA 9500 imager (GE Health Sciences).

### MD Simulations

Molecular dynamics (MD) simulations were performed for Rev1 and DNA complexes in explicit water solvent. Model preparation and simulations were performed using the AMBER v16 suite of programs for biomolecular simulations (31). AMBER’s *ff14SB* (32) force-fields were used for all simulations. MD simulations were performed using NVIDIA graphical processing units (GPUs) and AMBER’s *pmemd*.*cuda* simulation engine using our lab protocols published previously (33, 34).

A total of 3 separate simulations were performed (for binary, ternary and product complexes) based on the X-ray crystal structures determined in this study. The missing hydrogen atoms were added by AMBER’s *tleap* program. After processing the coordinates of the protein and substrate, all systems were neutralized by addition of counter-ions and the resulting system were solvated in a rectangular box of SPC/E water, with a 10 Å minimum distance between the protein and the edge of the periodic box. The prepared systems were equilibrated using a protocol described previously (35). The equilibrated systems were then used to run 1.0 μs of production MD under constant energy conditions (NVE ensemble). The use of NVE ensemble is preferred as it offers better computational stability and performance (36). The production simulations were performed at a temperature of 300 K. As NVE ensemble was used for production runs, these values correspond to initial temperature at start of simulations. Temperature adjusting thermostat was not used in simulations; over the course of 1.0 μs simulations the temperature fluctuated around 300 K with RMS fluctuations between 2-4 K, which is typical for well equilibrated systems. A total of 1,000 conformational snapshots (stored every 1,000 ps) collected for each system was used for analysis.

#### RMSF10 calculations

Root mean square fluctuations (RMSF) were computed based on the conformational snapshots collected during the MD simulations. To identify global motions on slower time-scales from MD, for each of the 3 systems the fluctuations associated with the first (slowest) 10 quasi-harmonic modes (RMSF_10_) were also computed and aggregated. It is well known that slowest 10 modes contribute to the majority of fluctuations in proteins (>80%) and the use of RMSF_10_, instead of all modes (RMSF), removes the faster stochastic motions of the protein, allowing focus on intrinsic dynamics of proteins (37). Both these calculations were performed using AMBER’s *ptraj* analysis program. All trajectory conformations were first aligned to a common structure, to remove any translation and overall molecular rotation during the simulations.

#### Protein-substrate interactions

The energy for the Rev1-DNA interactions (*E*_*Rev1-DNA*_) were calculated as a sum of electrostatic (*E*_*el*_) and van der Waals energy (*E*_*vdw*_) between atom pairs, based on an approach developed in previous publications (38, 39). All Rev1 and DNA atom pairs were included in the calculations and resulting interaction energies were summed up per residue pair. The energies were calculated for 1,000 snapshots, every 1,000 ps, sampled during the full 1.0 μs simulation and were averaged over these 1,000 snapshots.

## Acknowledgments

We thank Jay Nix (Molecular Biology Consortium 4.2.2 beamline at Advanced Light Source) for aid in remote data collection and help with data analysis. This research used resources of the Advanced Light Source, which is a Department of Energy Office of Science user facility under Contract DE-AC02-05CH11231. We thank Amy Whitaker (University of Kansas Medical Center) for helpful discussion and assistance with the manuscript preparation. This research was supported by the National Institute of General Medical Science [R35-GM128562 to B.D.F., T.M.W, L.M.C, T.H.K.] and [R01 - GM081433 to M.T.W].

---

**Supplemental Figure 1.**
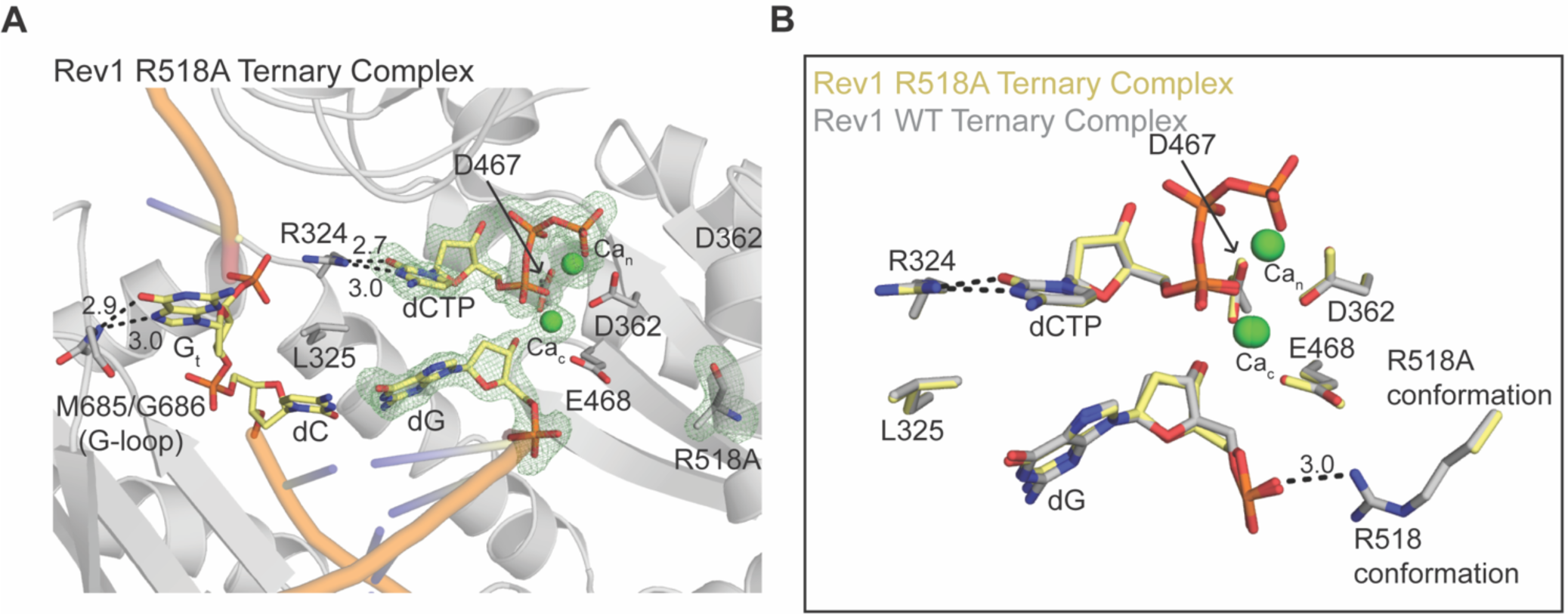
R518A Rev1 crystal structure. **(A)** An active site closeup of the R518A Rev1 crystal structure. The nucleic acid residues are shown in yellow and Rev1 in grey. An Fo-Fc map contoured at σ=3.0 around the incoming dCTP and primer terminal dG is shown as a green mesh. (**B)** An overlay of the R518A Rev1 binary (grey sticks) and ternary (yellow sticks) complexes are shown. Key protein and DNA residues are indicated in each panel.

